# Loss of estrogen unleashing neuro-inflammation increases the risk of Alzheimer’s disease in women

**DOI:** 10.1101/2022.09.19.508592

**Authors:** Fuhai Li, Inez Oh, Sayantan Kumar, Abdallah Eteleeb, Aditi Gupta, William Buchser, Chengjie Xiong, Sessions F. Cole, Eric McDade, Celeste M. Karch, Oscar Harari, Philip R. Payne, Carlos Cruchaga

## Abstract

The risk of Alzheimer’s disease (AD) in women is about 2 times greater than in men. The estrogen hypothesis is being accepted as the essential sex factor causing the sex difference in AD. Also, the recent meta-analysis using large-scale medical records data indicated estrogen replacement therapy. However, the underlying molecular targets and mechanisms explaining this sex difference in AD disease development remain unclear. In this study, we identified that estrogen treatment can strongly inhibition of neuro-inflammation signaling targets, using the systems pharmacology model; and identified ESR1/ESR2 (the receptors of estrogen) are topologically close to the neuroinflammation biomarker genes using signaling network analysis. Moreover, the estrogen level in women decreased to an extremely lower level than in men after age 55. Pooling together the multiple pieces of evidence, it is concluded that the loss of estrogen unleashing neuro-inflammation increases the women’s risk of Alzheimer’s disease. These analysis results provide novel supporting evidence explaining the potential mechanism of the anti-neuroinflammation role of estrogen causing the sex difference of AD. Medications boosting the direct downstream signaling of ESR1/ESR2, or inhibiting upstream signaling targets of neuroinflammation, like JAK2 inhibitors, on the signaling network can be potentially effective or synergistic combined with estrogen for AD prevention and treatment.

## Introduction

In 2022, it is estimated that about 6.5 million people are living with Alzheimer’s disease (AD), and the number of AD patients is predicted to rise to about 13 million^1^ in the United States. The estimated related healthcare cost is about 321 billion in 2022, which will increase to 1,000,000 million ($1 trillion)^1^. However, there is no effective and cure treatment yet. Recently, aducanumab is the only drug approved with limited effects for AD since 2003, though there are about 112 agents in the AD drug development pipeline by January, 2018^2^. These investigational agents target diverse AD-related molecular targets/signaling pathways and symptoms^2^. The failures of these AD treatment clinical trials are likely, in part, to limited knowledge of key molecular targets and complex signaling pathways e.g., oxidative stress, inflammation and immune response. Therefore, it is important to investigate the underlying molecular signaling mechanisms in females and males separately as novel therapeutic targets for AD treatment.

Sex is an important factor in AD. Among the individuals with AD, almost two-thirds are women, which means the risk of AD in women is about 2 times greater than in men^1,3^. In addition, AD is the 5^th^ leading cause of death for women (6.1% of deaths), and it is 2.6% of deaths for men; and more severe AD phenotypes in women than in men and faster memory decline^3,4,5,6^. The sex difference in AD indicated that female is one major risk factor for late-onset AD. Among the mechanisms explaining the sex difference in AD^7^, the estrogen hypothesis, which is neuro-protective, is being accepted as the sex factor causing the sex difference^8,9,10,11^. The effects of estrogen therapy for AD in clinical trial results are controversial and failed to show the beneficial effects^8^. Whereas, the recent large-scale meta-analysis indicated that estrogen replacement therapy (ERT) significantly decreased the risk of late-onset AD, with an odds ratio (OR) = 0.672, and Confidence interval (CI) = [0.581, 0.779]^12^.

On the other hand, neuro-inflammation has been identified as the third core feature of AD in recent years, in addition to Aβ and neurofibrillary tangles (NFTs). A few inflammation/immune genes are implicated, like TNF^13^, IL-1beta^14,15^, IL-6^15^, NFkB^16,17,18^. In the recent study^19^, a set of weakly activated neuro-inflammation and immune response signaling pathways were identified using bulk RNA-seq dataset of AD samples available from ROSMAP^20,21^ and Mayo Clinic^22^ datasets via systems biology and network analysis^19^. The uncovered neuroinflammation signaling targets are helpful to understand the dysfunctional neuroinflammation pathways. Studies have reported that one of the roles of estrogen is an anti-inflammation sex factor, and can be neuroprotection^10,11,23^. Whereas, the molecular targets and mechanisms of how estrogen inhibits neuroinflammation remain unclear.

In this study, using computational analysis, we identified that estrogen treatment can inhibit the uncovered neuroinflammation signaling targets; and inferred a signaling network that indicates the potential signaling pathways between estrogen receptors and the neuroinflammation signaling targets. On the other hand, using the life-span estrogen (female hormone) and testosterone (male hormone) level data in women and in men, it is found that in women, the estradiol hormone level (not the testosterone hormone) dramatically decreases to an extremely low starting from 45 years of age to 55 years of age. Whereas, in men, both estrogen and testosterone, stay relatively stable over the whole lifespan. Combing the two types of evidence, we concluded that loss of estrogen (not testosterone) unleashes neuro-inflammation and thus increases the risk of Alzheimer’s disease in women (not in men). Specifically, the three messages are: 1) estrogen can inhibit the uncovered neuron inflammation and immune response signaling; 2) ESR1 and ESR2, estrogen receptors, intensively interact with the neuroinflammation signaling; and 3) in women, estrogen decreased to an extremely low level (much lower than male) after around 55 years of age. Therefore, medications boosting the direct down-stream signaling of ESR1/ESR2, or inhibiting up-stream signaling of neuroinflammation on the signaling network can be potentially effective or synergistic combined with estrogen for AD prevention and treatment. The details of the methodology and results are presented in the following sections.

## Materials and Methodology

### Biomarker genes of the weakly activated inflammation and immune response signaling

In the recent study^19^, two sets of biomarker genes of the weakly activated neuroinflammation were identified using bulk RNA-seq dataset of AD samples available from ROSMAP^20,21^ and Mayo Clinic^22^ datasets. Specifically, 77 normal tissue samples and 81 AD tissue samples in Mayo Clinic dataset; and 260 normal samples and 97 AD samples in the ROSMAP dataset were used. By comparing the AD cases vs control, the intersect of weakly up-regulated genes in the two datasets was identified (with fold change >= 1.1; and p-value <= 0.1, which is defined as weakly activated molecular targets). By applying enrichment analysis on the KEGG signaling pathways, two major categories of signaling pathways were identified, i.e., inflammatory/immune response-related signaling pathways (like *Escherichia coli infection, pertussis, legionellosis, yersinia infection, malaria, NOD-like and IL-17 signaling*); and x-core signaling pathways (*like MAPK, NFkB, HIF-1, WNT, PI3K-AKT, Rap1, Hippo, TNF signaling pathways*) that topological closely interact with neuroinflammation on the KEGG signaling pathway network. In the inflammatory/immune response category signaling pathways, there are 121 up-regulated biomarker genes (see **Table 1**). In the x-core signaling pathways, there are 139 up-regulated biomarker genes (see **Table 2**). For more data analysis details, please refer to the study^19^. These two sets of biomarker genes will be used as gene set signatures to mine and rank drugs that can potentially inhibit the two sets of biomarker genes using the following connectivity map (CMAP) database ^24,25^.

**Table 1:**
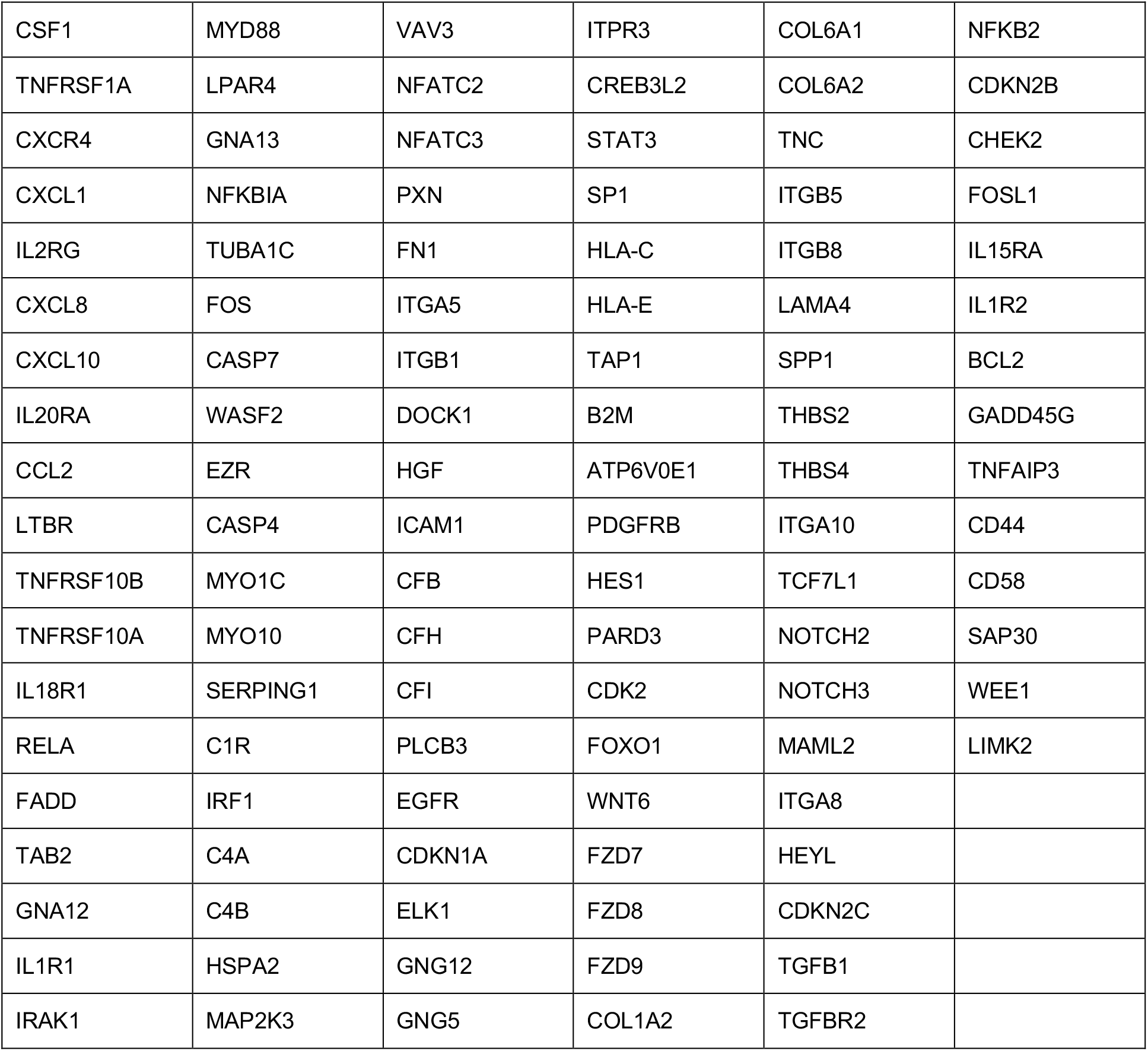
Biomarker genes of neuron inflammation and immune response.

**Table 2:**
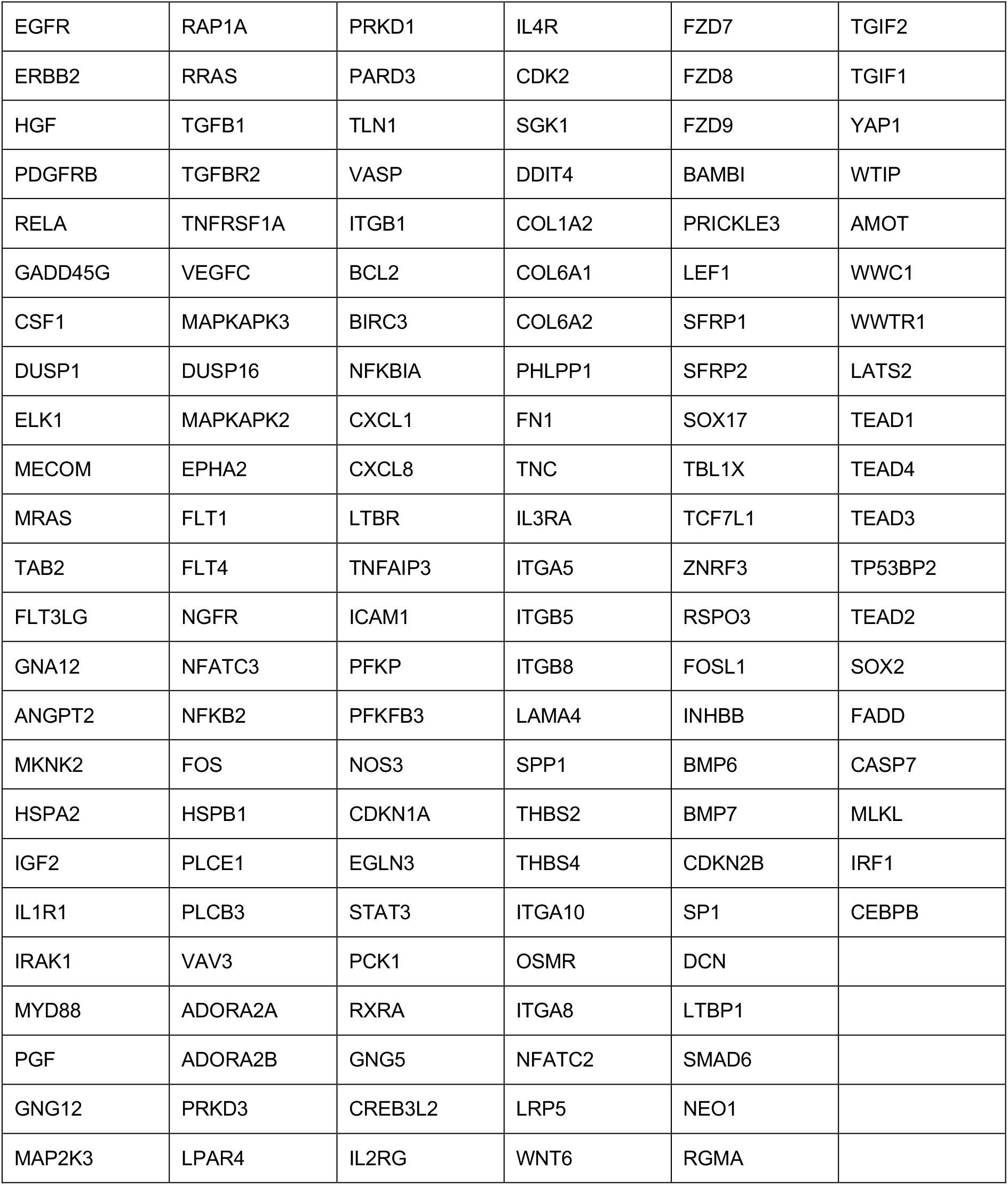
Biomarker genes of 9-core signaling pathways related to neuron inflammation and immune response.

### Mine drugs inhibiting the uncovered neuroinflammation signaling targets using the Connectivity Map (CMAP) and the two biomarker gene sets

Two sets of biomarker genes were used as the gene set signatures to identify potentially effective drugs that can inhibit the neuro-inflammation and immune signaling network biomarkers in the connectivity map (CMAP) database^24,25^. Specifically, the gene set enrichment analysis (GSEA)^26^ was applied on the gene set signatures respectively, and the z-profiles (gene expression variation before and after treatment with 2,513 drugs and investigational agents/compounds in the CMAP database) of nine (9) cancer cell lines to identify gene sets signature-specific inhibitory drugs. For each biomarker gene set, using the average GSEA scores across the 9 cancer cell lines, the drugs were ranked accordingly in ascending order. Then two sets of drugs and compounds/investigational agents that have a score <= -85 (indicating that these drug compounds could inhibit the activated biomarker genes) were selected respectively. Then the two drug lists were merged. Then, the selected drugs were annotated as different categories based on their targets and category information available from the CMAP database.

### ESR1/ESR2 and inflammation signaling network inference

The signaling pathways formatted in the protein-protein interaction (PPI) form in KEGG signaling pathway were collected. There were approximately 5,191 genes and 59,242 interactions. Then we conducted an unbiased and meaningful network inference to understand the potential signaling mechanism of how the neuroinflammation signaling targets interact with each other or other neighboring proteins on the KEGG signaling network. Specifically, the average shortest path distance of all other genes on the KEGG signaling network (other than the union of the two sets of biomarker genes) was calculated. Then the neighbor genes that are close to the biomarker genes, i.e., with an average distance <= 2.8 (empirically selected), were selected. These unbiasedly selected neighbor genes indicated the potential targets intensively interacting with the neuroinflammation biomarker genes. The identified neuroinflammation signaling network indicated the potential biomarkers that can regulate and perturb the activated neuron inflammation and immune response signaling targets.

### Identify FDA-approved drugs inhibiting genes on the uncovered signaling network

The FDA-approved drug and their target information were derived from DrugBank (version 5.0) database^27^. Then the FDA-approved drugs inhibiting the selected genes were identified as drug candidates that can potentially perturb the uncovered signaling network.

### Electronic healthcare records-based analysis of the effect of estrogen use on dementia progression

Structured electronic healthcare records (EHR) data spanning the timeframe from June 1 2008 to May 31 2018 were extracted from the Washington University School of Medicine/Barnes-Jewish Hospital Allscripts TouchWorks database for patients seen the Memory Diagnostic Center at Washington University School of Medicine in St. Louis. These data were obtained and analyzed with approval from the Washington University Institutional Review Board. Dementia progression was tracked by looking at longitudinal changes in global Clinical Dementia Rating^®^ (CDR^®^) scores: 0 denotes normal cognition, 0.5 very mild dementia, 1 mild dementia, 2 moderate dementia, and 3 severe dementia. Estrogen use was identified from the medications table via string searching of medication names and RxNorm code matching. Male patients and patients who had only fewer than two global CDR scores were excluded from the analysis. Patients were stratified by estrogen use and the average change in CDR score over time for the estrogen-using group versus the non-estrogen-using group was compared using a Wilcoxon rank sum test. These data were obtained and analyzed with approval from the Washington University Institutional Review Board under protocol #<u>201905161</u>.

## Results

### Two sets of neuro-inflammation related biomarkers genes

As aforementioned in the method section, in the recent study^19^, two (2) sets of neuroinflammation related biomarker genes were identified, as seen in the following **Table 1** (109 biomarker genes in the inflammatory/immune response related signaling pathways) and **Table 2** (139 biomarker genes in the x-core signaling pathways) respectively. The two sets of biomarker genes were then used as gene set signatures to mine and rank drugs respectively that can potentially inhibit the two sets of biomarker genes using the following connectivity map (CMAP) database ^24,25^.

### Estrogen can inhibit the two sets of gene biomarkers

For each of the two sets of biomarker genes as seen in **Table 1** and **Table 2**, drugs and compounds that have an average GSEA score <= -85 (indicating that these drug compounds could inhibit the activated gene set signatures in each biomarker gene set) were selected respectively, and then merged together. To understand the top-ranked/selected drugs, we clustered these drugs/compounds, and annotate the drug clusters as different categories based on their targets and category information available from the CMAP database, as seen in **Fig. 1**. Interestingly, the estrogen receptor agonist category was identified as the potential effective anti-neuroinflammation treatment. Also there are a set of drugs with diverse mechanisms or modes of action (MOA) were identified. As seen in the top part of the figure, immune suppression, including the JAK2 inhibitors, and anti-inflammatory drugs/compounds categories were top-ranked. Also, there are multiple categories related to fatty acid oxidation, fatty acide synthase, lipoxygenase. It is known that dysfunctional lipid signaling occurred in AD. On the right of **Fig. 1**, angiogenesis inhibitor, VEGFR, EGFR, FGFR, PDGFR tyrosine, RAF, KIT inhibitors were top-ranked. It indicated the dysfunctional or broken angiogenesis functions in AD. From another view point, it might indicate that the brain tries to generate new blood vessels to repair the damaged angiogenesis by secreting these growth factors. Also, the adenylate cyclase activator is related to the activation of cAMP which can potentially activate autophagy signaling. The calcium channel blocker is associated with the dysfunctional calcium (Ca2+) signaling in AD. In the bottom-right part of **Fig. 1**, the TNF inhibitor, dipeptidyl peptidase inhibitor (for type2 diabetes (T2D) treatment) and HMGCR inhibitors (statins), and NFkB pathway inhibitor, HIV integrase inhibitor, and nitric oxide production inhibitor were identified. In addition to the categories, there are a set of solo category drugs, like adrenergic receptor, aetylcholine receptor, and tyrosine phosphatase inhibitors. Also, the TGFbeta, MEK, JNK, MAPK, p38 and MTOR inhibitors were also top-ranked.

**Figure 1:**
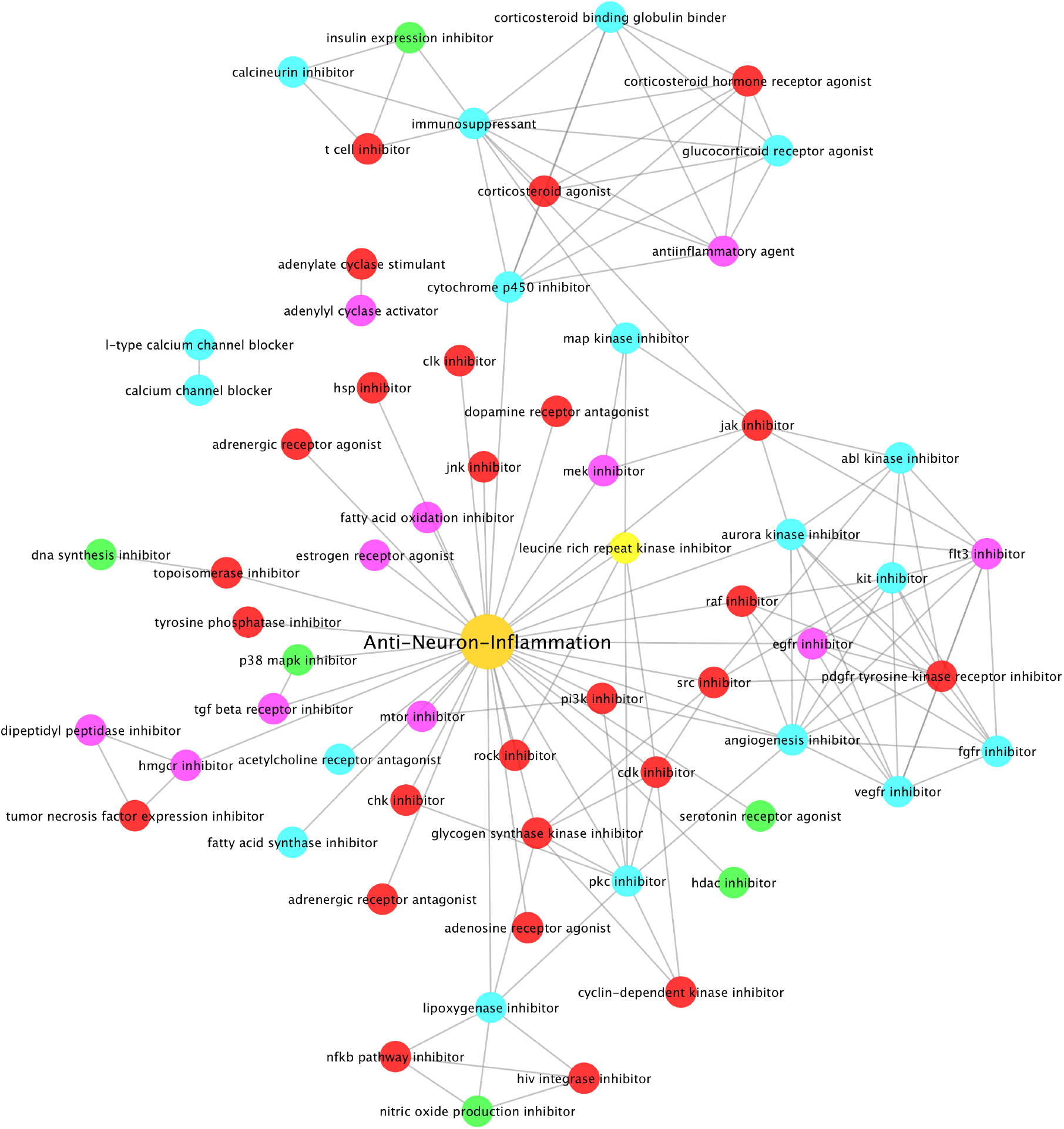
Top-ranked drug categories that can potentially inhibit activated neuron inflammation and immune response. The connection (edge) between two different categories indicates that some drugs belong to both categories. Color was randomly assigned for visualization purpose.

To further investigate the specific drugs/compounds of these drug categories, we mapped the drugs to the selected drug categories, as seen in **Fig. 2**. As seen, the estrogen hormones, including estriol, estrone, estradiol, diethylstilbestrol and propyl pyrazole triol (PPT) in the estrogen receptor agonist category were identified to be able to inhibit the neuron inflammation signaling. The immunosuppressant, anti-inflammation, corticosteroid agonist and glucocorticoid receptor agonist, like the drug dexamethasone (a NR3C1 inhibitor), were also top-ranked. Moreover, the JAK inhibitors, like tofacitinib, lestaurtinib, tozasertib were top-ranked. Interestingly, the dexamethasone and JAK2 inhibitors, are the first and second reported drugs to be able to significantly reduce the mortality rate of COVID-19 patients validated in clinical trials^28^,^29^. Thus, drugs from different categories with potentially different modes of action (MOA), and their combinations/cocktails (targeting multi-signaling modules/dysfunctional signaling pathways in AD) can be potentially effective and synergistic to inhibit neuron inflammation for AD treatment and prevention.

**Figure 2:**
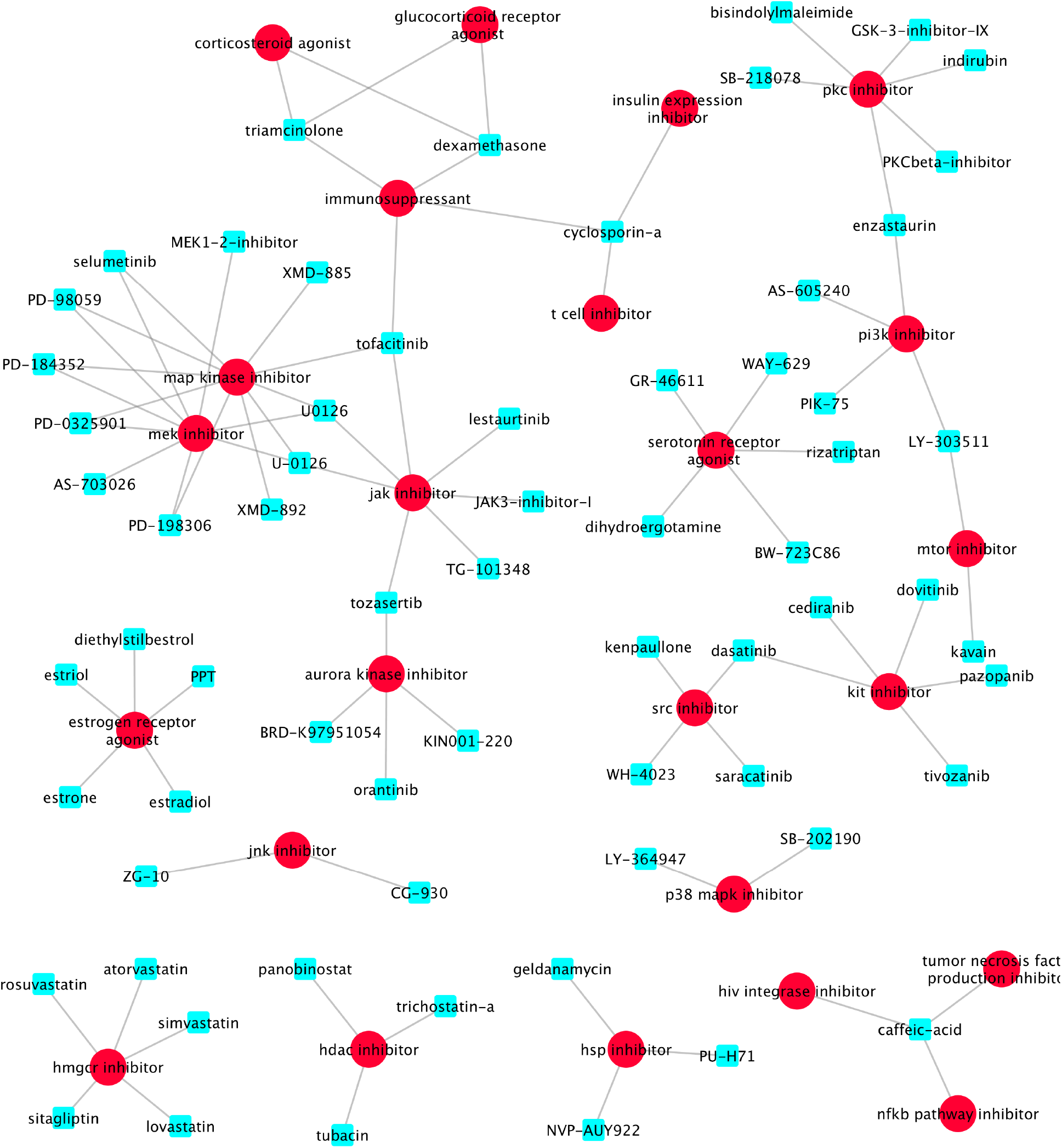
Top-ranked drugs of some selected drug categories.

### ESR1 and ESR2, receptors of estrogen, are topologically close to the neuroinflammation biomarker genes on the KEGG signaling network

It is known that ESR1 and ESR2 are estrogen receptors. Therefore, it is interesting to know that if ESR1/ESR2 plays important roles in the anti-neuroinflammation of estrogen. In another word, it is interesting and critical to understanding the potential consequent signaling pathways of estrogen because the consequent signaling pathways can provide mechanistic insights of how estrogen inhibits neuro-inflammation; and can identify additional therapeutic targets for managing neuroinflammation. As aforementioned in the network inference part in the Method section, we identified the potential signaling network that is topologically close to these neuroinflammation-related biomarker genes (the union of the two sets of biomarker genes). Specifically, the average distance between inflammation genes and these neighbor genes is <= 2.8 connection path steps (number of signaling interactions) in the corresponding shortest paths on the KEGG signaling pathways. **Fig. 3** shows the neuron inflammation and immune response signaling network linking the 361 genes with 4,922 signaling interactions. The cyan color nodes indicated the neighbor genes that are close to the uncovered neuron inflammation and immune response biomarker gene sets. Interestingly, ESR1 and ESR2, the receptors of estrogen, appeared as the neighbor genes of the neuroinflammation signaling targets. It indicated that the ESR1 and ESR2 are close to and intensively interact with the neuron inflammation genes. Thus, it serves as independent evidence of the anti-neuroinflammation role of estrogen potentially via ESR1 and ESR2.

**Figure 3:**
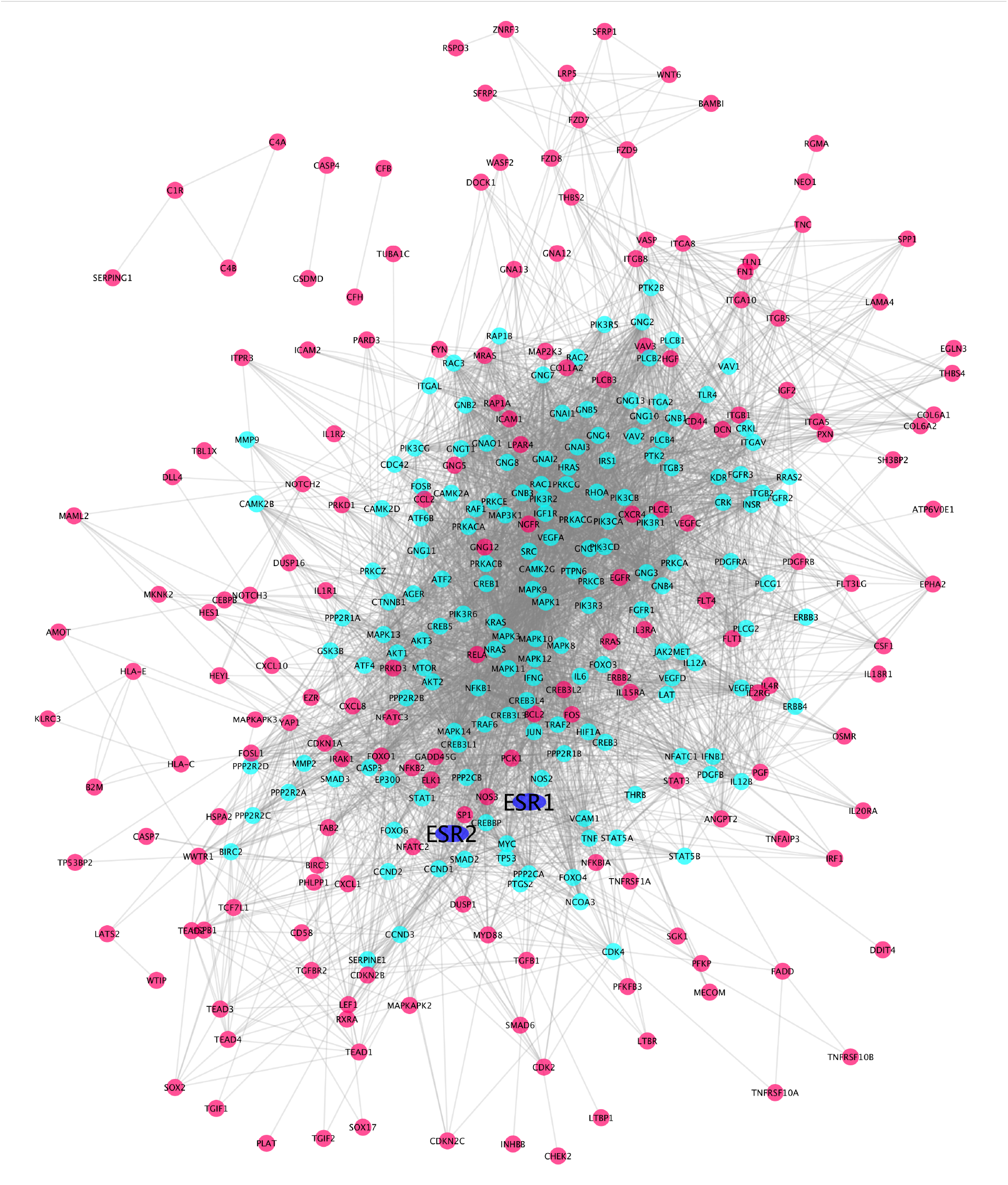
Neuron inflammation and immune response signaling network (with 4,922 interactions among 361 proteins). Red: neuroinflammation related biomarker genes. Cyan: genes topologically close to the neuroinflammation biomarker genes. Blue: ESR1 and ESR2 (the known two estrogen receptors).

Moreover, as seen in **Fig. 3**, it is interesting that there is a set of signaling targets in the central part of the signaling network surrounded by neuroinflammation signaling targets. These signaling targets in the center of the uncovered signaling network, like the *MAPK, JAK2, STAT, PI3K, SRC, CREB, ATK, and MYC* genes, could be additionally effective therapeutic targets of anti-neuroinflammation. Moreover, the molecular mechanisms of how the ESR1 and ESR2 interact with these targets to inhibit neuroinflammation could provide novel therapeutic targets for discovering effective drugs and drug combinations for AD treatment and prevention. In another word, medications boosting the direct down-stream signaling of ESR1/ESR2, or inhibiting up-stream signaling of neuroinflammation, like JAK2 inhibitors, on the signaling network can be potentially effective or synergistic combined with estrogen for AD prevention and treatment.

### Estrogen decreases to an extremely low level in women around the age of 55

If estrogen has the anti-neuroinflammation effect, why do women have a double risk probability to have AD, and have a more severe disease phenotype than men, considering women have a much higher level of estrogen? To resolve the question, we collected and analyzed the life-span change data of estrogen in women and men from the website of Centers for Disease Control and Prevention (CDC)^23, 30^. **Fig. 4-upper panel** shows the average life-span changes of estrogen in women and men. As seen, in women, estrogen decreases starting from about the age of 45; reached the same level as in men; and dramatically decreased to an extremely low level (close to zero (0), and much lower than in men) around 55 years of age. Whereas the estrogen level in men is constantly maintained. Therefore, the difference in life-span estrogen level change in females and males can explain why women have double the risk probability to have AD, and have more severe disease phenotype than men. Before the onset of AD, the disease begins about 20 or more years, and 73% of AD patients are diagnosed at age 75 or older^1,31^. The loss of estrogen to an extremely low level is correlated with the beginning of AD in women.

**Figure 4:**
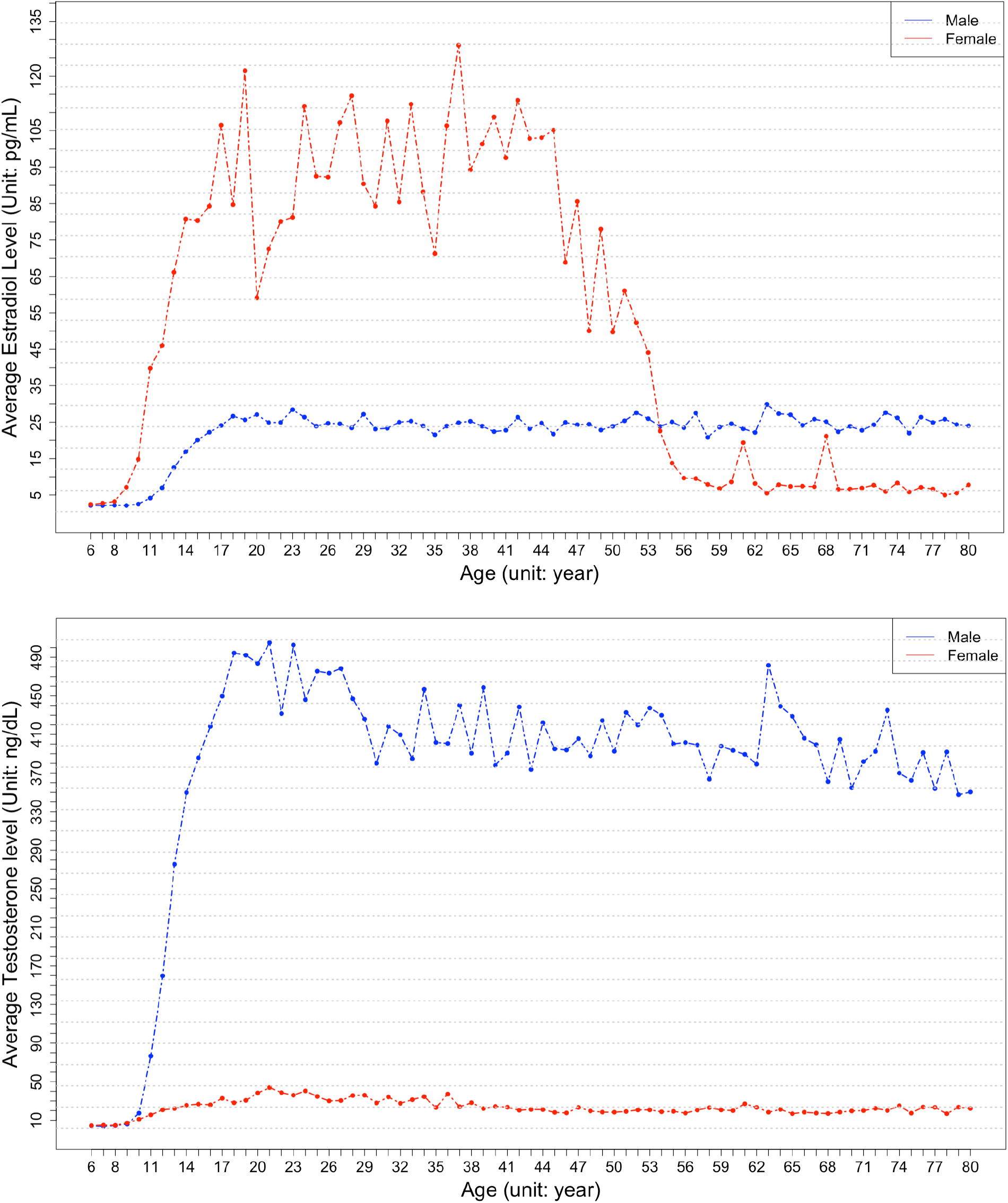
The lifespan changes of estradiol (female sex hormone, **upper-panel**) and testosterone (male sex hormone, **lower-panel**) level in female (red) and male (blue). As seen, in women, the estradiol hormone level (not the testosterone hormone) dramatically decreases to an extremely low starting from 45 years of age to 55 years of age. Whereas, in men, both estrogen and testosterone, stay relatively stable over the whole lifespan.

The male sex hormone, androgen has been associated with anti-inflammation effects^32,33^. The life-span change of testosterone (male sex hormone - androgen) data was also collected from CDC, as seen in **Fig. 4-lower panel**. Different from estrogen, the testosterone level in males decreases slowly and keeps at a high level for the whole lifespan in men. Also, the testosterone level stays stable in females. As shown in **Fig. 4**, interestingly, young men roughly have about 1/4 (25 pg/ml vs 100 pg/ml) level of estrogen in young women; whereas, young women have only 1/10 (40 ng/dl vs 400 ng/dl) level of androgen in young men. It indicated that estrogen has stronger anti-inflammation effects than androgen. Thus, estrogen, as an anti-inflammation factor, is needed for both men and women; whereas, androgen is mainly useful for men. In conclusion, the loss of estrogen to almost zero level (rather than androgen) unleashes neuroinflammation, and subsequently increases the risk of Alzheimer’s disease in women (not in men).

### Longitudinal analysis of electronic healthcare records data indicated that estrogen treatment can slow down Clinical Dementia Rating (CDR) score progression in dementia patients

We analyzed electronic healthcare records data to investigate the relationship between estrogen treatment and the Clinical Dementia Rating (CDR), which is a standard score to indicate dementia severity, with a higher CDR indicating more severe dementia. Global CDR was found to increase at a slower rate in the patients who used estrogen (n = 71) than the patients who did not use estrogen (n = 1,004) (0.194 versus 0.298 CDR increase per year, *p* = 0.045). This suggests that estrogen treatment has a protective effect on dementia progression.

## 4. Discussion and Conclusions

There are about 2 times more women AD patients than men, and female patients develop AD phenotypes faster than male patients. Estrogen is believed as an essential sex factor causing the sex difference in AD. Moreover, the recent meta-analysis using large-scale medical records data indicated estrogen replacement therapy. However, the underlying molecular mechanisms explaining the effects of estrogen in AD disease development remain unclear. In this study, we showed 3 types of novel evidence to explain the potential mechanism of the anti-neuroinflammation role of estrogen causing the sex difference of AD: 1) estrogen treatment can inhibit neuro-inflammation signaling targets; 2) ESR1/ESR2 (the receptors of estrogen) are topologically close to the neuroinflammation biomarker genes; and 3) estrogen level in women decreased to an extremely lower level (almost to 0) after age 55. Pooling together the multiple pieces of evidence, it is concluded that the loss of estrogen unleashing neuro-inflammation increases the women’s risk of Alzheimer’s disease. Therefore, estrogen and additional medications boosting the direct down-stream signaling of ESR1/ESR2, or directly inhibiting neuroinflammation, can be synergistic cocktails for AD prevention and treatment.

There are some limitations to our work, which require further attention. The detailed mechanism of neuroinflammation remains unclear. Microglia are the immune cells in brain, and play important roles in neuroinflammation regulation. It is important to evaluate the effects of estrogen on different cells in the brain, like neurons, microglia, and astrocytes. Using the single nuclei/cell RNA-seq of AD patients with and without estrogen treatment might be useful to investigate the effects of estrogen treatment on different cell types. As shown in **Fig. 4**, estrogen should have a stronger anti-inflammation capability, which is important for both men and women. Thus, it is interesting to investigate the optimal estrogen level, like the average level in men, for the purpose of anti-neuroinflammation. Though the meta-analysis indicated the effectiveness of estrogen treatment in AD, it did not show significant improvement in AD treatment and prevention in clinical trials, which might be due to the late-stage AD is not reversible. It is interesting to investigate the effects of estrogen in early-stage AD. Moreover, as indicated in the signaling network analysis, the up-stream signaling pathways of neuroinflammation; or the down-stream signaling pathways of neuroinflammation could be additional therapeutic targets for AD treatment and prevention. The current version of a network linking the neuroinflammation and ESR1/ESR2 can be further improved by combining computational network analysis with the estrogen perturbations on different cell types, to uncover the fine-scale signaling targets and pathways in different cell types. Since there is no effective treatment for AD. Drug cocktails/combinations targeting multiple essential signaling targets; or targeting multi-cell types could be novel and effective regimens for AD treatment and prevention.

## Acknowledgment

This work is partially supported by the National Institute of Ageing (NIA) R56AG065352 to Dr. Li. We thank all the participants and their families, as well as the many involved institutions and their staff. Funding: This work was supported by grants from the National Institutes of Health (R01AG044546 (CC), P01AG003991(CC), RF1AG053303 (CC), RF1AG058501 (CC), and U01AG058922 (CC), and chuck Zuckerberg initiative (CZI). This work was supported by access to equipment made possible by the Hope Center for Neurological Disorders, and the Departments of Neurology and Psychiatry at Washington University School of Medicine. CC receives research support from Biogen, EISAI, Alector and Parabon. CC is a member of the advisory board of Vivid Genomics, Halia Therapeutics and ADx Healthcare. The remaining authors declare no competing interests.

